# Multidimensional plasticity in the Glanville fritillary butterfly: larval performance curves are temperature, host and family specific

**DOI:** 10.1101/2020.05.05.065698

**Authors:** Nadja Verspagen, Suvi Ikonen, Marjo Saastamoinen, Erik van Bergen

**Affiliations:** Helsinki Institute of Life Science, University of Helsinki, Finland; Research Centre of Ecological Change, Faculty of Biological and Environmental Sciences, University of Helsinki, Finland; Lammi Biological Station, University of Helsinki, Finland

**Keywords:** developmental plasticity, GxExE, intraspecific variation, temperature, nutrition, multidimensional plasticity

## Abstract

Variation in environmental conditions during development can lead to changes in life-history traits with long-lasting effects. Here, we study environmentally induced variation, i.e. the consequences of potential maternal oviposition choices, in a suite of life-history traits in pre-diapause larvae of the Glanville fritillary butterfly. We focus on offspring survival, early growth rates and relative fat reserves, and pay specific attention to intraspecific variation in the responses (GxExE). Globally, we found that thermal performance and survival curves varied between diets of two host plants, suggesting that host modifies the temperature impact, or *vice versa*. Additionally, we show that the relative fat content has a host-dependent, discontinuous response to developmental temperature. This implies that a potential switch in resource allocation, from more investment in growth at lower temperatures to storage at higher temperatures, is dependent on other environmental variables. Interestingly, we find that a large proportion of the variance in larval performance is explained by differences among families, or interactions with this variable. Finally, we demonstrate that these family-specific responses to the host plant remain largely consistent across thermal environments. Altogether, the results of our study underscore the importance of paying attention to intraspecific trait variation in the field of evolutionary ecology.

## 1. Introduction

Species can cope with environmental change by avoiding stressful conditions, by producing phenotypes better adjusted to the new environmental conditions through plasticity, or by adapting to the novel conditions through evolutionary change [1, 2]. Even though the avoidance of environmental stress is an effective strategy, e.g. through tracking favourable conditions by expanding to higher latitudes or altitudes [3], it is often limited by factors such as the distribution of resources, the structure of the landscape and/or the dispersal ability of the species. Moreover, when environmental changes are rapid, adaptive evolution may not occur fast enough. In those cases, plasticity can enable species to persist under the novel conditions, allowing more time for mutations to arise and selection to occur [4, 5]. Assessing a species’ ability to respond plastically to environmental change, and evaluating its performance when exposed to conditions that are beyond or at the limit of the normal range, could therefore shed light on whether organisms will be able to persist future conditions.

Developmental plasticity is defined as the process through which external conditions, such as nutrition and temperature, can influence developmental trajectories and lead to irreversible changes in the adult phenotype [1]. This phenomenon is ubiquitous in nature, especially among taxa that have sessile life-styles [6-8]. The environmental regulation of development has been studied extensively using insects, whose pre-adult stages are often immobile and thus must cope with local environmental conditions. In general, when exposed to higher temperatures, insect larvae tend to grow faster [9, 10] and the body size of the emerging adults is smaller [9, 11, 12], which might alter performance later in life. Likewise, nutrition is known to regulate development in insects through nutrient balance [13-15] and/or the concentration of secondary metabolites in the diet [16].

When assessing responses to changes in environmental conditions, it is important to recognise that the environmental factors that affect the phenotype typically occur simultaneously and interactively [17]. Hence, plastic responses to one type of environmental stress may be dependent on the state of another external factor. Such non-additive multidimensional plasticity, in response to combinations of thermal and nutritional environments, has been demonstrated in moths [14], butterflies [18] and fruit flies [19]. For example, Singh et al. showed that poor host plant quality mainly influenced development at intermediate temperatures the tropical butterfly *Bicyclus anynana* [18]. Moreover, significant genetic variation for (multidimensional) plasticity is known to exist in both natural and laboratory populations [20-22]. This intraspecific variation in the ability to respond to an environmental cue (GxE), or combinations of cues (GxExE), is hypothesised to be beneficial in the light of climate change since it facilitates the evolution of wider ranges of environmental tolerance [23, 24].

In this study we focus on environmentally induced variation in a suite of life-history traits in the Glanville fritillary butterfly (*Melitaea cinxia*). The species occurs at its northern range margin on the Åland archipelago (SW Finland) where it inhabits a highly fragmented network of habitat patches that are defined by the presence of at least one of two available host plant species; *Plantago lanceolata* and *Veronica spicata*, hereafter referred to as *Plantago* and *Veronica*, respectively [25]. Adult females produce large egg clutches, and the selection of suitable oviposition sites is known to be a hierarchical process [26, 27]. In the field, gravid females of the Glanville fritillary appear to first choose habitats that are hot, dry and sunny [28, 29]. Host plant discrimination, with individuals typically preferring one host species over the other, occurs subsequently [30, 31]. Therefore, selective mothers can influence the developmental trajectories of their offspring through oviposition site selection, which in turn may affect offspring performance and fitness [32].

Using a full-factorial split-brood design, we explore the consequences of these maternal oviposition choices for the pre-diapause larvae of *Melitaea cinxia*. We aim to research the (combined) effects of developmental temperature and host plant on pre-diapause larvae of this species. We measure the survival, early growth rates and relative fat content of offspring reared at four different temperatures and on two different host plants, and pay attention to intraspecific variation in the responses by using individuals from different genetic backgrounds (i.e. families). As shown in other insects, we expect a large positive effect of developmental temperature on growth rate. Furthermore, in the scenario of additive multidimensional plasticity, we expect larvae to grow faster and have higher survival on *Veronica* within each thermal environment, as this has previously been demonstrated under laboratory conditions [16, 33]. Individuals that develop fast, and thus will be diapausing for longer, are predicted to allocate relatively more resources to fat storage, which is thought to be the primary fuel for overwintering and post-winter activities. Finally, given that the natural habitat of this species is heterogeneous and fragmented, we expect family-specific responses to the environmental factors (GxE, GxExE) to be important determinants of the phenotype.

## 2. Methods

### Study system

*Melitaea cinxia* is a univoltine species and on the Åland islands adults emerge from their pupae in June, after which females lay several clutches of 100 – 200 eggs [34]. The sessile pre-diapause larvae hatch in late June and early July and live gregariously on the host plant of their mothers’ choice. In the beginning of autumn the larvae spin a communal web in which they diapause until spring. Overwinter survival is impacted by multiple factors, among which body size, with larger larvae having a higher chance to survive [35]. After diapause, larvae become solitary and can move over longer distances in search of resources and/or suitable microhabitats [36]. The laboratory population of *M. cinxia* used in this study was established in 2015 from 136 post-diapause larvae (consisting of 105 unique families) collected from 34 habitat patches across the large network of habitat patches on the Åland islands.

### Experimental design

In the spring of 2019, diapausing larvae of the laboratory stock were stimulated to recommence development, reared to adulthood in small transparent plastic containers, and mated with an unrelated individual. Subsequently, the gravid females were provided with a single *Plantago* plant for oviposition, and the host plant was checked daily for newly produced egg clutches. Clutches were carefully removed, placed in individual petri-dishes, and transferred to a climate‐controlled cabinet set to 28:15 °C and a 12L:12D cycle.

Egg clutches of 15 females were divided over eight experimental treatments in two steps, yielding a full-factorial split-brood design with two diets of different host plants (*Plantago* and *Veronica*) and four developmental day temperatures (28 °C, 30 °C, 32 °C and 34 °C). First, to ensure the utilization of a single host plant species throughout development, egg clutches were divided into two equal groups 3-5 days after oviposition. One of these groups was provided with fresh leaves of *Plantago* while the other received fresh *Veronica* leaves. All plants were reared under standard conditions (28:15 °C). Second, when approximately 90% of the larvae within each group transitioned from the first to the second instar, we generated experimental cohorts of 15 siblings. These cohorts were randomly divided over four climate-controlled chambers (28:15 °C, 30:15 °C, 32:15 °C and 34:15 °C, all with a 12L:12D cycle, and using a Sanyo MLR-350 for the 32 °C treatments and a Sanyo MLR-351 for the others).

Throughout the experiment, larvae were inspected every morning and fresh leaves were provided to ensure *ad libitum* feeding conditions. For five families, individuals from a second clutch (from the same parents) were used to complete all experimental treatments. One female did not produce enough offspring to complete all treatments and these data have been excluded from further analyses. A schematic representation of the experimental design is given in Figure S1. For further information on the background of larvae used in the experiment see Table S1.

### Life-history traits

We studied environmentally induced variation in a suite of life-history traits and focussed on offspring survival, early growth rates, and the relative amount of fat reserves accumulated during early development. To assess offspring survival, the larvae within each cohort were counted every fourth day, and on these days the entire cohort was also weighed to the nearest 0.01 mg (Mettler Toledo XS105 DualRange) to trace overall mass gain during early development. This procedure was continued until the first individual of the cohort entered the diapause stage, which can be recognised by a change in body colour (from pale-brown to black), an increase in larval body hair density, and the presence of red eyes. From this date forward individual data was collected by recording the day of entering diapause and the body mass of each diapausing larvae. Subsequently, larvae were frozen to -80 °C, and stored in eppendorf tubes until further processing.

The individual growth rates were calculated according to the formula

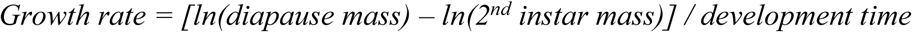

where 2^nd^ instar mass (i.e. mass at the start of the experiment) was estimated by dividing the mass of the entire cohort by the number of individuals, and development time was computed as the time between the start of the experiment and the day the individual entered diapause [37].

Relative fat content at diapause was determined for seven randomly chosen individuals per cohort. These larvae were dried to constant mass (60 °C for 24 h) and weighed to the nearest 0.01 mg, yielding initial dry mass. Triglyceride and free fatty acids were extracted by incubating the dried body at room temperature in 1:2 (v/v) methanol:dichloromethane for 72 h, followed by drying and re-weighing, yielding fat-free dry mass [38]. The relative fat content was calculated according to the formula

*Relative fat content = (initial dry mass - fat-free dry mass) / initial dry mass*

### Statistical analyses

Interval-censored survival curves were fitted using the *survival* package [39] and plotted using the *survminer* package [40]. Log-rank tests were performed to determine the influence of temperature and host plant on survival using the *interval* package [41]. A linear model was fitted to estimate the effect of temperature and host on the mean amount of body mass gained during early development. Cohort mass was divided by the number of surviving individuals and log-transformed to improve normality. The day of the experiment, temperature and host plant (and all interactions) were included in the full model. Two additional linear models were fitted to estimate the effect of family, temperature and host (and all interactions) on individual growth rate and relative fat content. For all models described above, step-wise model selection based on AIC values was performed using the *step()* function. Post hoc pairwise comparisons (Tukey’s HSD; α = 0.05) were performed using the *emmeans* package [42].

Intraspecific variation in the responses to the host plant is explored by extracting the slope of a linear model – with individual growth rate as dependent and host plant as independent variable – for each family and within each thermal environment. These slopes describe both the magnitude and the direction of the response to the host plant [20]. Using Pearson correlations we test whether host-induced responses (i.e. the slopes) are family-specific and consistent across thermal environments. All statistical analyses were performed in R [43].

## 3. Results

### Pre-diapause survival and clutch mass

Probability of survival was generally high but dropped considerably for larvae with long development times (i.e. those that enter diapause in the upper percentiles of the distribution of development times, Figure 1). The probability of survival was not affected by temperature (asymptotic logrank two-sample t-test, P = 0.3968 for individuals reared on *Plantago*, and P = 0.8678 for individuals reared on *Veronica*). Survival was significantly lower for larvae that were reared on *Plantago*, but only for the two highest temperatures (P-values given in Figure 1).

**Figure 1.**
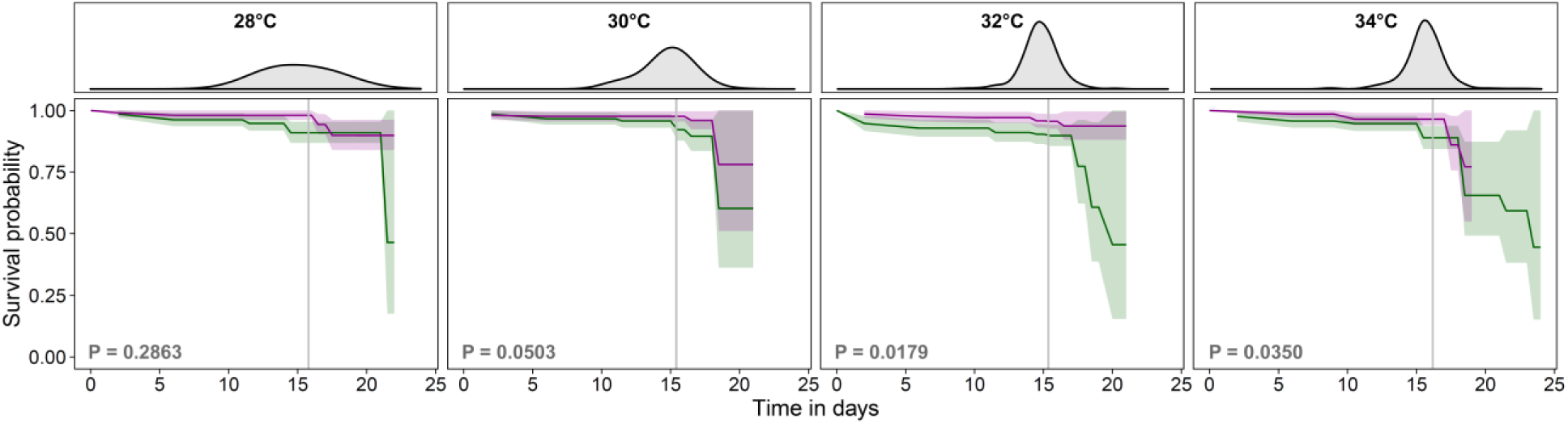
Kaplan-Meier survival probability over time for larvae reared on *Plantago* (green) and *Veronica* (purple), at four different day temperatures. Shaded area represents the 95% confidence interval. Grey lines show the mean day of diapause, and the distribution of diapausing day is given in the upper panels. The probability of survival is not affected by temperature, but, at the two highest day temperatures, survival is significantly lower for larvae that were reared on *Plantago* (P-values given in the figure).

We found that both the thermal environment and the host plant interacted with time to affect mean clutch mass. The effect of temperature on mean clutch mass increases with time (time:temperature, F = 10.5182, P < 0.001), with larvae reared at 28 °C being significantly smaller than those reared at higher temperatures from day 8 onward (Figure 2A). The mean clutch mass of cohorts reared on *Veronica* increased faster over time compared to those reared on *Plantago* (time:plant, F = 3.9190, P = 0.0089). Cohorts using *Veronica* were smaller than those using *Plantago* at the start of the experiment (day 0; Figure 2A) but larger at the final time point (day 16).

**Figure 2.**
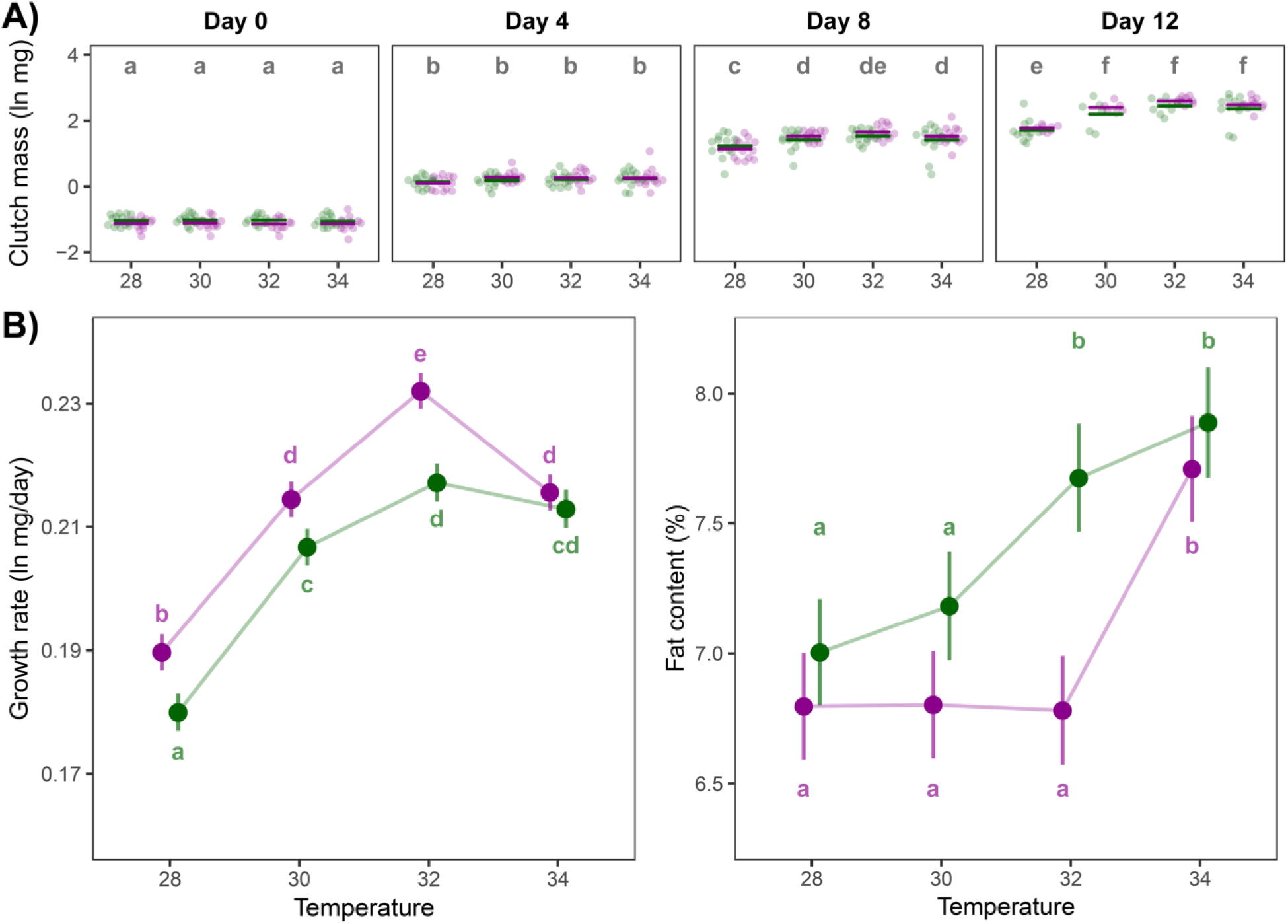
Environmentally induced variation in life-history traits. **A)** Dots depict the mean clutch mass, log-transformed and corrected for number of individuals, in each thermal environment (x-axis), for *Plantago* (green) and *Veronica* (purple), on four assessment days (from left to right: day 0 [i.e. 2^nd^ instar mass], day 4, day 8 and day 12). Significant differences between thermal treatments (Tukey’s HSD, *α =* 0.05) are indicated by different letters. Details of the statistical test can be found in Table S2. **B)** Model-estimated marginal means for the individual growth rates (left; R^2^ = 0.5964) and the relative fat content (right; R^2^ = 0.4992). Error bars represent 95% confidence intervals and significant differences between groups (Tukey’s HSD, α = 0.05), averaged over the families, are indicated by different letters. Details of statistical tests can be found in Tables S3 and S4.

### Individual growth rates and allocation to fat reserves

For both life-history traits (growth rate and fat content) we found that all main effects and all interaction terms were statistically significant (see Tables S3 and S4). Averaged over the families, model-estimated marginal means for the individual growth rates revealed that individuals achieve higher growth rates on *Veronica*, except for those reared at 34 °C (Figure 2B, Table S3C). Growth rate increased with temperature until a maximum at 32 °C. At an even higher temperature of 34 °C growth rate dropped significantly compared to that at 32 °C for larvae fed with *Veronica* (pairwise comparison: P < 0.001). In contrast, the growth rates of larvae reared at 34 °C on *Plantago* were not significantly different from those of individuals reared at 32 °C (pairwise comparison: P = 0.5213). This decrease in growth rate at 34 °C was mainly caused by an increase in development time rather than a decrease in body mass (Figure S2).

The relative fat content showed a discontinuous change to the temperature gradient on both hosts (Figure 2B, Table S4C). For individuals reared on *Plantago*, development at the two higher temperatures resulted in significantly higher relative amounts of fat reserves. The thermal threshold at which change in relative fat content occurs was higher for individuals reared on *Veronica*, where only development at the highest temperature lead to an increase in relative fat content. As a result of the difference in threshold we only observed a significant effect of host plant at 32 °C (pairwise comparison: P < 0.001), with larvae utilizing *Plantago* having a higher relative fat content on average.

### Family-specific responses to the host plant

Our results demonstrate that intraspecific variation for multidimensional phenotypic plasticity (GxExE) is large in this system. For both life-history traits, but especially for the individual growth rates, the (interactive) effects of environmental cues were highly dissimilar across families. About 12% of the total phenotypic variance (V_P_) in individual growth rates was explained by the interaction between the family and the host plant (family:host plant, F = 32.2507, P < 0.001; see table S3B). In other words, family-specific responses to the host were an important determinant of the phenotype. Indeed, some families used in the experiment achieved the highest growth rates on *Veronica* while individuals from other families grew consistently faster on *Plantago* (Figure 3). These family-specific reaction norm slopes were positively correlated across thermal environments (*Pearson*’s *r* 0.4-0.8; Figure S3). Moreover, utilising *Plantago* as a host plant resulted in higher variance in larval growth rates across families (and not within families; Figure S4).

**Figure 3.**
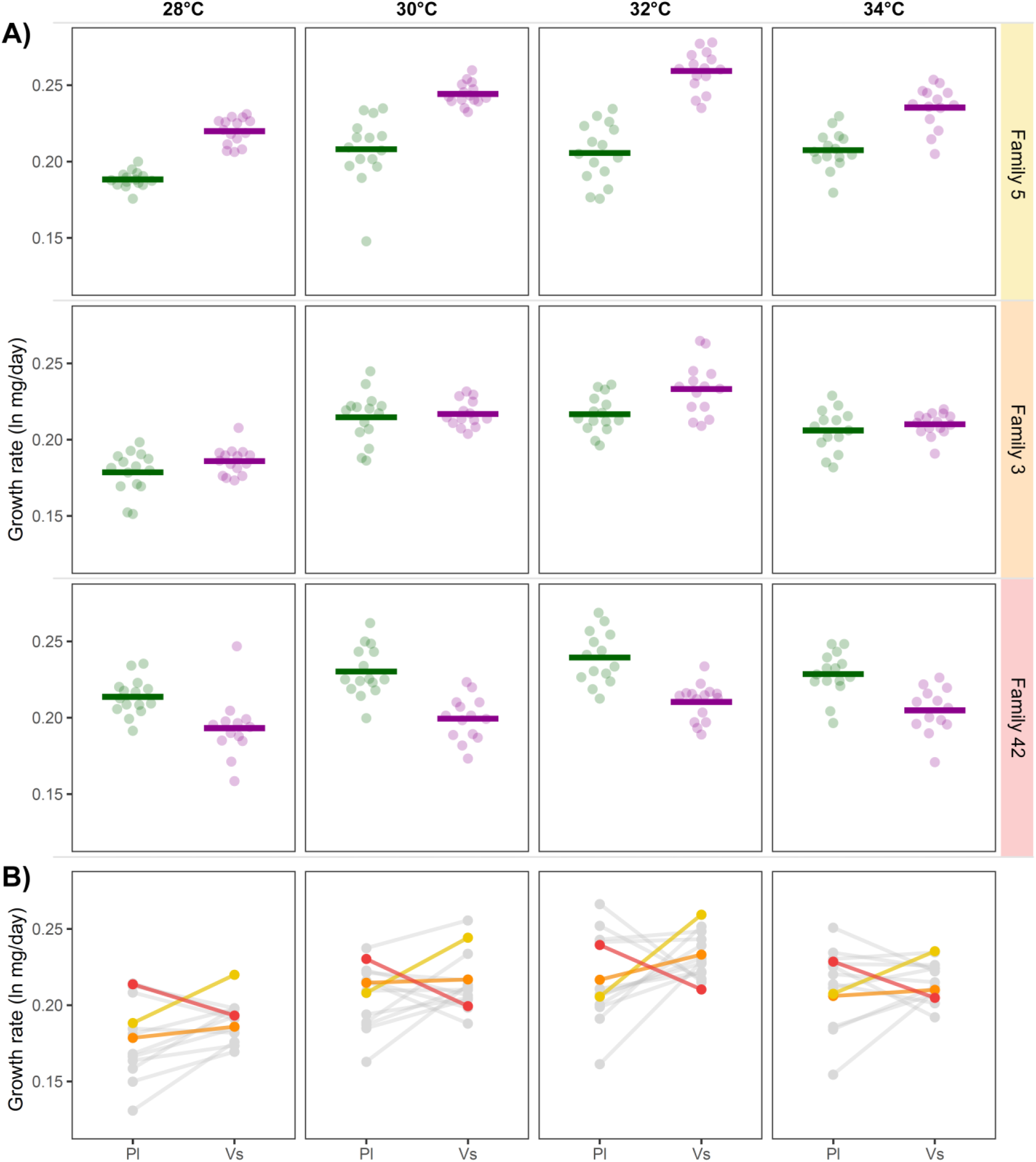
Host-induced responses in growth rates vary across families, but are consistent across thermal environments. **A)** Panels demonstrate the individual growth rates on *Plantago* and *Veronica* of three representative families. Siblings of some families consistently achieve higher growth rates on *Veronica* (upper panels; yellow), while individuals from other families demonstrate an equal performance on each host plant (central panels; orange) or even grow faster on *Plantago* (lower panels; red). **B)** Norms of reaction to the host plant for all families included in the experiment, coloured families correspond to those given in panel A. The reaction norm slopes, describing both the magnitude and the direction of the response, are family-specific and correlate strongly across thermal environments (for details see Figure S3A). Utilising *Plantago* leads to higher between-family variance in growth rates (for details see Figure S3B).

## 4. Discussion

Using a full factorial design, with fourteen genetic backgrounds (families), four developmental temperatures and two host plant species, we explored the relative contributions of different sources of phenotypic variance across a suite of life-history traits in the Glanville fritillary butterfly. We start this section by describing the general patterns observed in our data, and then discuss how the developmental trajectories of pre-diapause larvae could be influenced by maternal oviposition site selection in the wild. Subsequently, we go into the variation in environmental responses observed among families in our study, and discuss how this genetic variation for (multidimensional) plasticity may impact the population’s ability to persist environmental heterogeneity.

In ectotherms, temperature can affect developmental processes directly through changes in chemical reaction kinematics and the physical properties of membranes [44, 45], which in turn can impact organismal performance and fitness. Some developing individuals are able to manipulate their thermal environment, for example by relocating to warmer microhabitats. Alternatively, when immature life-stages are largely immobile, such as in the case of the Glanville fritillary butterfly, the optimal thermal environment for development can be realized through selective oviposition choices of the female. As is true for many butterfly species, Glanville fritillary mothers could regulate the thermal environment of their offspring by preferring sunny or shady environments for oviposition [28, 29]. Averaged over the families, our data showed a clear initial increase in growth rate with increasing temperature, with an optimum around 32 °C. At higher temperatures the growth rate decreased for larvae reared on *Veronica* and stabilised for those reared on *Plantago*.

Maternal preferences to oviposit in sunny habitats, thereby increasing the developmental temperatures of their offspring, are therefore intuitively considered to be adaptive. However, even though the average ambient day temperatures on Åland are well below 32 °C during the end of summer, when pre-diapause larvae are developing (Figure S5), sunshine creates thermal stratification which can cause the temperatures close to the ground to rise to be between 12 and 20 °C above ambient temperature [46, 47]. This suggests that the maternal preference for sunny habitat could be maladaptive when the ambient day temperatures on Åland are above 20 °C (see also Salgado, 2020). In this scenario developmental temperatures in sunny microclimates may rise well above the observed optimal temperature of 32 °C for larval growth, and potentially even exceed the thermal tolerance limits of the larvae. It was recently shown that the summer of 2018 was an anomaly in terms of precipitation, temperature and vegetation productivity across the habitats of the Glanville fritillary butterfly in Åland, and that this extreme climatic event was associated with a 10-fold demographic decline of the metapopulation [48]. Though this dramatic decline has been attributed to severe water deficits during May and July, the record-breaking temperatures observed for July (above 25 °C; figure S5) could have exceeded the thermal tolerance limits of pre-diapause larvae developing in sunny environments.

In addition to an effect of temperature on growth rates, our data also reveal a general trend in the relative fat content of the larvae. This physiological trait is important for butterfly life- history[49] and was quantified for the first time in this species. Relative fat content increases significantly between 30 and 32 °C for larvae reared on *Plantago*, and between 32 and 34 °C for larvae reared on *Veronica*. Previous research has shown similar increases in relative fat content with increasing temperature in other insects [e.g. 50, 51, 52], while other studies have described the opposite pattern [e.g. 38, 53]. Our hypothesis, stating that individuals predicted to spend more time in diapause, i.e. those with shorter development times, accumulate more fat during development was therefore falsified. In fact, the cohorts with the largest relative fat reserves also demonstrated the longest development times and thus shortest time in diapause (i.e. those reared at 34 °C). Instead, variation in relative fat content seemed to be associated with the relative investment in growth. Individuals reared at 32 °C allocated more resources to their fat reserves when they utilized *Plantago* instead of *Veronica*. At this temperature we also observed the largest difference in host-specific growth rates, with individuals utilizing *Plantago* demonstrating reduced investment in growth compared to their siblings reared on *Veronica.* This suggests that individuals of this species trade-off growth for increased investment in fat storage at higher temperatures, and that the thermal threshold for this switch in life history strategies is influenced by the larval diet.

Within the preferred sunny habitats, mothers select one of two available host plants which may differ in their suitability for larval development. Overall, and in concordance with our hypothesis and earlier reports [16, 33], we found that pre-diapause larvae perform better on *Veronica* than on *Plantago*. Though survival in the first instars was high on both plant species, the probability of survival to diapause was higher for larvae that were reared on *Veronica*, particularly in the two hottest environments. In addition, individuals on this host achieved higher average growth rates in most thermal environments. Finally, the larval performances of the families used in this study were very uniform when utilizing *Veronica*, which was in stark contrast to the high variance in growth rates observed among families on *Plantago*.

Since females can maximize their fitness by laying their eggs on host plants on which the performance of their offspring is maximized (preference–performance hypothesis), Glanville fritillary mothers are predicted to prefer *Veronica* when both hosts are available. Interestingly, it has been shown that females of this species do not necessarily prefer the host plant that is most abundant in their local environment, but that this preference depends on which host is more abundant at a larger regional scale [30, 54, 55]. This local adaptation is attributed to the spatial distribution of the two hosts in the field, with *Plantago* being omnipresent and *Veronica* mainly occurring in habitat patches in the north-western part of the archipelago. Females from regions where *Veronica* is an abundant and therefore reliable host plant were observed to prefer *Veronica* when offered a binary choice, while butterflies in regions where *Veronica* is less reliable preferred to oviposit on *Plantago* [30, 55].

It is important to place the observed effects of the host plant on larval performance in the context of our experimental design; while temperature interacted with the host plant at the level of the insect herbivore, direct effects of temperature on the host were not assessed. The plants used in the study were cultivated and kept under greenhouse conditions throughout the experiment, and thus represented a high quality diet for the developing larvae. In nature, the two host plants themselves may differ in their responses to variation in temperature, and this interspecific variation may affect the nutritional quality of the available hosts for example in terms of primary nutrients. Additionally, the two host plants used in this study both produce iridoid glycosides [56] — defence chemicals known to deter feeding by generalist insect herbivores [57, 58]. Interestingly, these iridoids can also act as feeding stimulants in specialist butterfly larvae [59, 60], such as *M. cinxia,* and the concentrations of these secondary compounds are known to be susceptible to variability in precipitation and temperature in *Plantago* [61]. Thus, in the field, when potential effects of temperature on the host plant are present, the interacting effects of temperature and host on insect development demonstrated here could be different.

In addition to the more general patterns described above, we found significant genetic variation for (multidimensional) plasticity in this system. In other words, the phenotypic responses to the (combination of) environmental variables were highly dissimilar across families (i.e. significant GxE and GxExE interactions). For example, the interaction between the family and the host plant explained a large proportion of the variance in individual growth rates, and these family-specific responses to the host were largely consistent cross thermal environments. While most families grew faster on *Veronica* (e.g. family 5), others consistently achieved their highest growth rates on *Plantago* (e.g. family 42). Such intraspecific variation, or in this case variation within the meta-population, for plasticity is common in both natural and laboratory populations [e.g. 20, 62] and hypothesised to be beneficial for insects exposed to climate change [63, 64] because it increases their evolutionary potential [24].

As a final note we would like to emphasize that the general patterns described in studies like the one presented here (i.e. using a relatively small number of families), may be susceptible to bias when phenotypes vary strongly across genetic backgrounds. For example, using fourteen families we describe that larvae of *M. cinxia* in general perform better on *Veronica* than on *Plantago*, while this is in fact not the case for all families. Therefore, with a different and/or smaller subset of families we could potentially have observed different general patterns.

In conclusion, we demonstrated that larval performance curves in the Glanville fritillary butterfly are family-specific and interactively mediated by the thermal and nutritional environment. The results of our study therefore underscore the importance of studying the multidimensionality of environmental effects on phenotype expression. In addition, our work demonstrates that intraspecific variation is likely an important determinant of population-level responses to environmental change in this system.

## Acknowledgements

We are grateful to Heini Karvinen for practical assistance during the experiments, and Wilco Verberk for suggestions that greatly improved earlier versions of our manuscript. Financial support was provided by European Research Council (independent starting grant no. 470 637412 ‘META-STRESS’ to MS). NV was supported by the Erasmus+ programme of the European Union and a Lammi Biological Station’s Environmental Research Foundation (LBAYS) grant.

## Authors’ Contributions

NV, SI, MS and EvB conceived and designed the experiments. NV performed the experiments. NV and EvB analysed the data and wrote the first draft of the manuscript. All authors provided critical feedback and helped to shape the research, analyses and manuscript.

## Data Accessibility

Raw data will be archived in the Dryad Digital Repository upon acceptance of the manuscript.

## Supplementary Materials

### Contents

Figure S1: Experimental design.

Figure S2: Mean development time and diapause mass per developmental temperature and host plant.

Figure S3: Family specific responses to host plant.

Figure S4: Minimum, mean and maximum temperatures in Åland.

Table S1: Background of larvae used in the experiment.

Table S2: Linear model for mean clutch mass.

Table S3: Linear model for individual growth rates.

Table S4: Linear model for individual fat content.

Table S5: Linear model for individual development time.

Table S6: Linear model for individual diapause mass.

**Figure S1:**
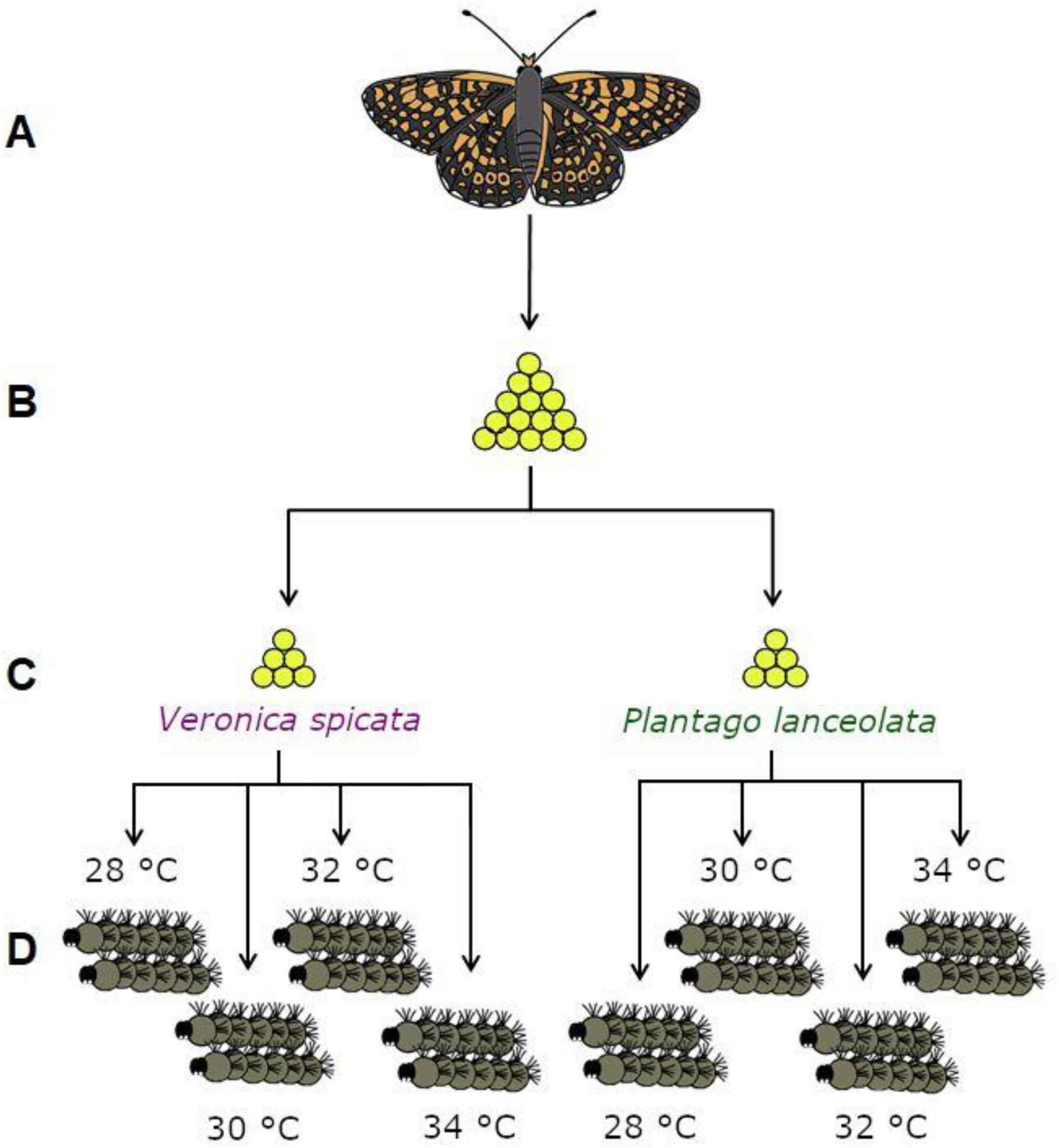
Schematic outline of the experimental design. The egg clutch of one female (A + B) was split over two host plants (C), and further divided over four temperature treatments upon transitioning to the second instar (15 larvae per treatment, D). This was done for offspring of 15 females from different families.

**Figure S2:**
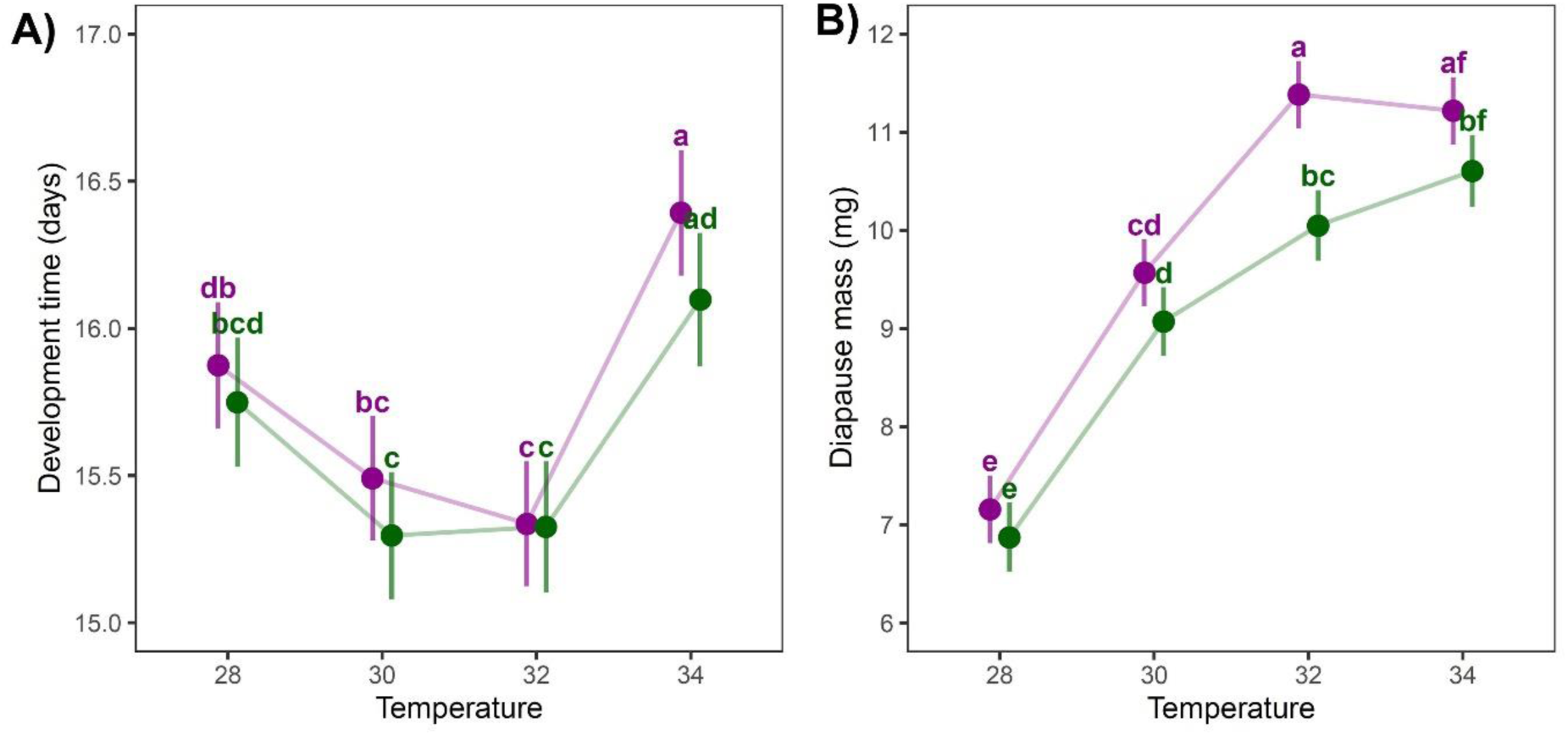
Environmentally induced variation in life history traits. Model-estimated marginal means for the individual **A**) development time (R^2^ = 0.3941) and **B**) diapause mass (R^2^ = 0.5283). Error bars represent 95% confidence intervals and significant differences between groups (Tukey’s HSD, α = 0.05), averaged over the families, are indicated by different letters. Details of statistical tests can be found in Tables S5 and S6.

**Figure S3:**
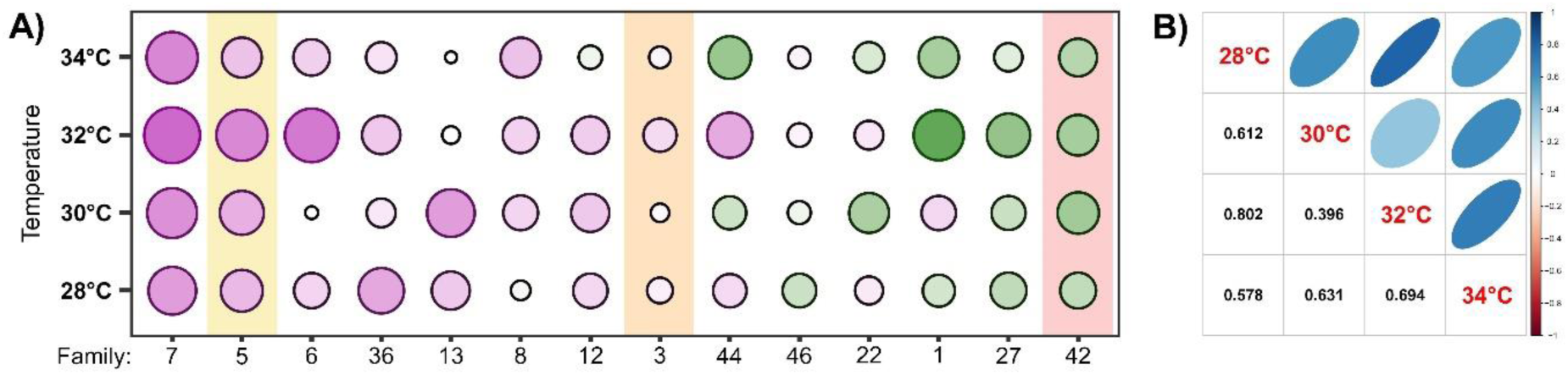
Family-specific responses to the host plant. **A**) Some families consistently achieve higher growth rates on *Veronica* while other families, regardless of the thermal environment, grow consistently faster on *Plantago.* The size of the symbol depicts the magnitude of the host-induced response, with steeper reaction norm slopes represented by larger symbols. The direction of the response to the host plant is represented by the colour of the symbol, with higher growth rates on *Veronica* depicted in purple and higher growth rates on *Plantago* in green. Highlighted families correspond to those given in figure 3A of the main text. **B**) Pearson’s correlation coefficients among the host-induced reaction norm slopes were positive and ranged between 0.4 and 0.8. The host-induced responses are therefore family-specific and largely consistent across thermal environments.

**Figure S4:**
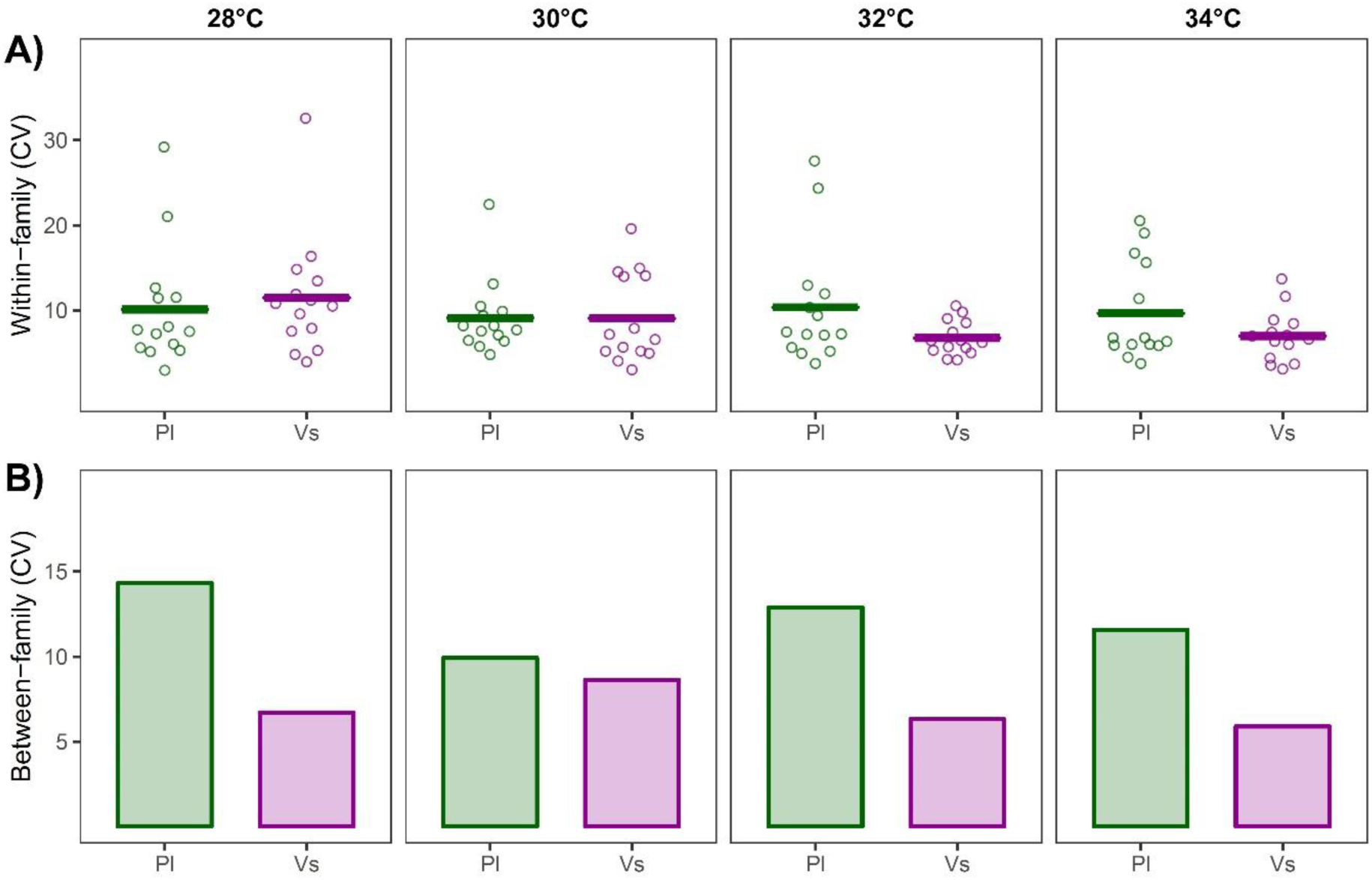
**A**) Variance in larval growth rates within families (CV; standard deviation divided by the mean for each family) was similar between host treatments. **B**) Utilising *Plantago* as a host plant resulted in higher variance in larval growth rates across families (CV; standard deviation of family means divided by the global mean growth rate, calculated for each temperature treatment separately)

**Figure S5:**
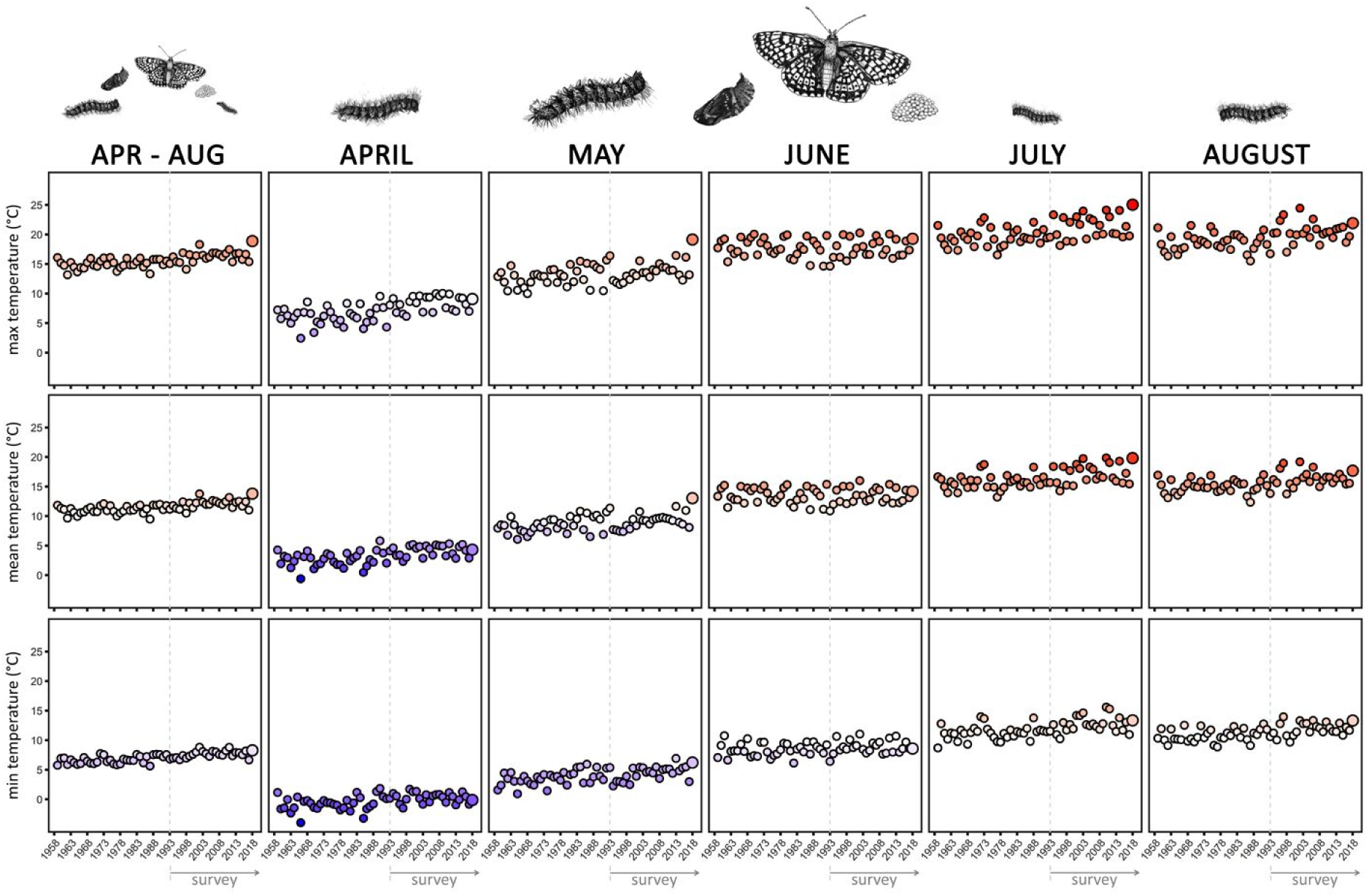
Minimum, mean and maximum temperatures in Åland in the months April until August for years between 1958 and 2018. Symbol colour gradient indicates temperature, with cold temperatures represented by blue and warm temperatures indicated by red. The pictures above the panels show the presence of butterfly life-stages per time period, with post diapause larvae being present in April and May, after which they pupate and emerge as adults who lay their eggs in June. Pre-diapause larvae then develop in July and August. Temperature data was derived from the Jomala climate station database in Åland. Illustrations courtesy of Luisa Woestmann.

**Table S1:**
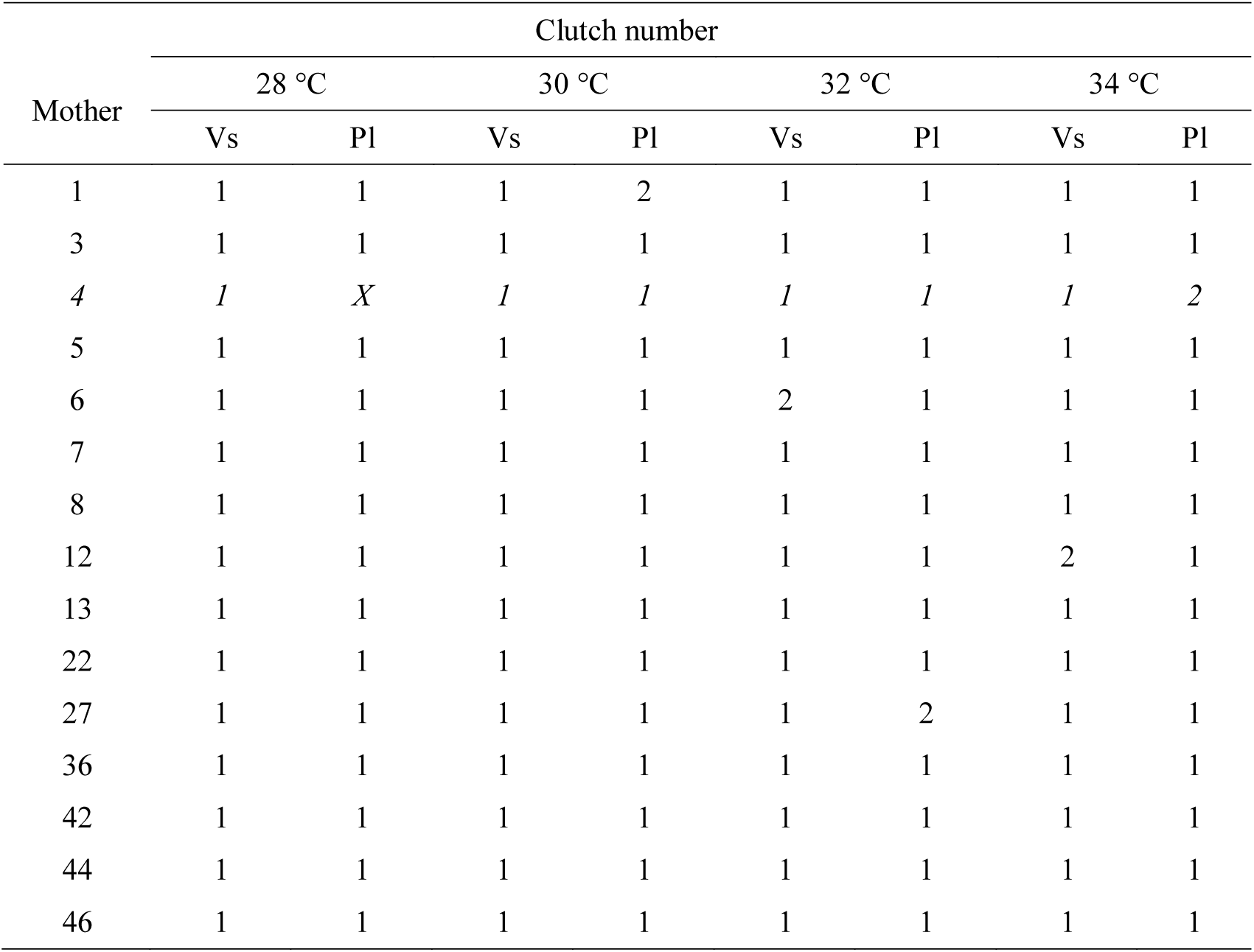
Background of larvae used in the experiment including family of the mother and the clutch number. Larvae printed in italics were not included in the data analysis. Abbreviation Vs stands for host plant *Veronica spicata*, Pl for host plant *Plantago lanceolata*.

| Mother | Clutch number |  |  |  |  |  |  |  |
| --- | --- | --- | --- | --- | --- | --- | --- | --- |
|  | 28 °C |  | 30 °C |  | 32 °C |  | 34 °C |  |
|  | Vs | Pl | Vs | Pl | Vs | Pl | Vs | Pl |
| 1 | 1 | 1 | 1 | 2 | 1 | 1 | 1 | 1 |
| 3 | 1 | 1 | 1 | 1 | 1 | 1 | 1 | 1 |
| 4 | *1* | *X* | *1* | *1* | *1* | *1* | *1* | 2 |
| 5 | 1 | 1 | 1 | 1 | 1 | 1 | 1 | 1 |
| 6 | 1 | 1 | 1 | 1 | 2 | 1 | 1 | 1 |
| 7 | 1 | 1 | 1 | 1 | 1 | 1 | 1 | 1 |
| 8 | 1 | 1 | 1 | 1 | 1 | 1 | 1 | 1 |
| 12 | 1 | 1 | 1 | 1 | 1 | 1 | 2 | 1 |
| 13 | 1 | 1 | 1 | 1 | 1 | 1 | 1 | 1 |
| 22 | 1 | 1 | 1 | 1 | 1 | 1 | 1 | 1 |
| 27 | 1 | 1 | 1 | 1 | 1 | 2 | 1 | 1 |
| 36 | 1 | 1 | 1 | 1 | 1 | 1 | 1 | 1 |
| 42 | 1 | 1 | 1 | 1 | 1 | 1 | 1 | 1 |
| 44 | 1 | 1 | 1 | 1 | 1 | 1 | 1 | 1 |
| 46 | 1 | 1 | 1 | 1 | 1 | 1 | 1 | 1 |

**Table S2:** Linear model for mean clutch mass (related to Figure 2A in the main text). **A**) Minimum adequate model (in bold) was obtained using the *step()* function, starting from the full model. **B**) Anova table for the minimum adequate model. **C**) Model-estimated marginal means, as well as upper and lower confidence limits, for all experimental treatments. Significant differences experimental groups (Tukey’s HSD, α = 0.05), averaged over the families and host plants, were obtained using the package *emmeans* and are indicated by different letters.

| **A** |  | AIC |
| --- | --- | --- |
| M<sub>full</sub> | Ln(mass) ~ t * T * HP | -1162.5 |
| M<sub>1</sub> | Ln(mass) ~ t + T + HP + t:T + t:HP + T:HP | -1177.6 |
| **M<sub>final</sub>** | **Ln(mass) ~ t + T + HP + t:T + t: HP** | **-1179.8** |
| M<sub>3</sub> | Ln(mass) ~ t + T + HP + t:T + T:HP | -1171.2 |
| M<sub>4</sub> | Ln(mass) ~ t + T + HP + t:HP + T:HP | -1107.7 |
| M<sub>5</sub> | Ln(mass) ~ t + T + HP + t:T | -1173.6 |
| M<sub>6</sub> | Ln(mass) ~ t + T + HP + t: HP | -1111.8 |

t = time-point; T = temperature; HP = host plant.
| **B** | Df | Sum Sq | Mean Sq | F value | P value | % exp |
| --- | --- | --- | --- | --- | --- | --- |
| Time-point | 3 | 615.2663 | 205.0888 | 3768.1423 | < 0.0001 | 94.9792 |
| Temperature | 3 | 5.2870 | 1.7623 | 32.3799 | < 0.0001 | 0.8162 |
| Plant | 1 | 0.1098 | 0.1098 | 2.0169 | 0.1564 | 0.0169 |
| Time-point:Temperature | 9 | 5.1522 | 0.5725 | 10.5182 | < 0.0001 | 0.7954 |
| Time-point:Plant | 3 | 0.6399 | 0.2133 | 3.9190 | 0.0089 | 0.0988 |
| Residuals | 392 | 21.3354 | 0.0544 |  |  | 3.2936 |

| **C** | Temperature | Time-point | Mean | LCL | UCL | Group |
| --- | --- | --- | --- | --- | --- | --- |
|  | 28 °C | 0 | -1.0789 | -1.16619 | -0.99152 | a |
|  | 30 °C | 0 | -1.0636 | -1.15095 | -0.97628 | a |
|  | 32 °C | 0 | -1.0795 | -1.16684 | -0.99217 | a |
|  | 34 °C | 0 | -1.0926 | -1.17996 | -1.00529 | a |
|  | 28 °C | 4 | 0.1221 | 0.03477 | 0.20944 | a |
|  | 30 °C | 4 | 0.2316 | 0.14423 | 0.31890 | a |
|  | 32 °C | 4 | 0.2416 | 0.15425 | 0.32891 | a |
|  | 34 °C | 4 | 0.2509 | 0.16352 | 0.33819 | a |
|  | 28 °C | 8 | 1.1837 | 1.09635 | 1.27102 | b |
|  | 30 °C | 8 | 1.4689 | 1.38152 | 1.55619 | bc |
|  | 32 °C | 8 | 1.5895 | 1.50220 | 1.67687 | cg |
|  | 34 °C | 8 | 1.4655 | 1.37820 | 1.55287 | cd |
|  | 28 °C | 12 | 1.7373 | 1.63179 | 1.84272 | g |
|  | 30 °C | 12 | 2.2997 | 2.16625 | 2.43306 | e |
|  | 32 °C | 12 | 2.5193 | 2.41542 | 2.62313 | fe |
|  | 34 °C | 12 | 2.4235 | 2.32922 | 2.51788 | e |

**Table S3:**
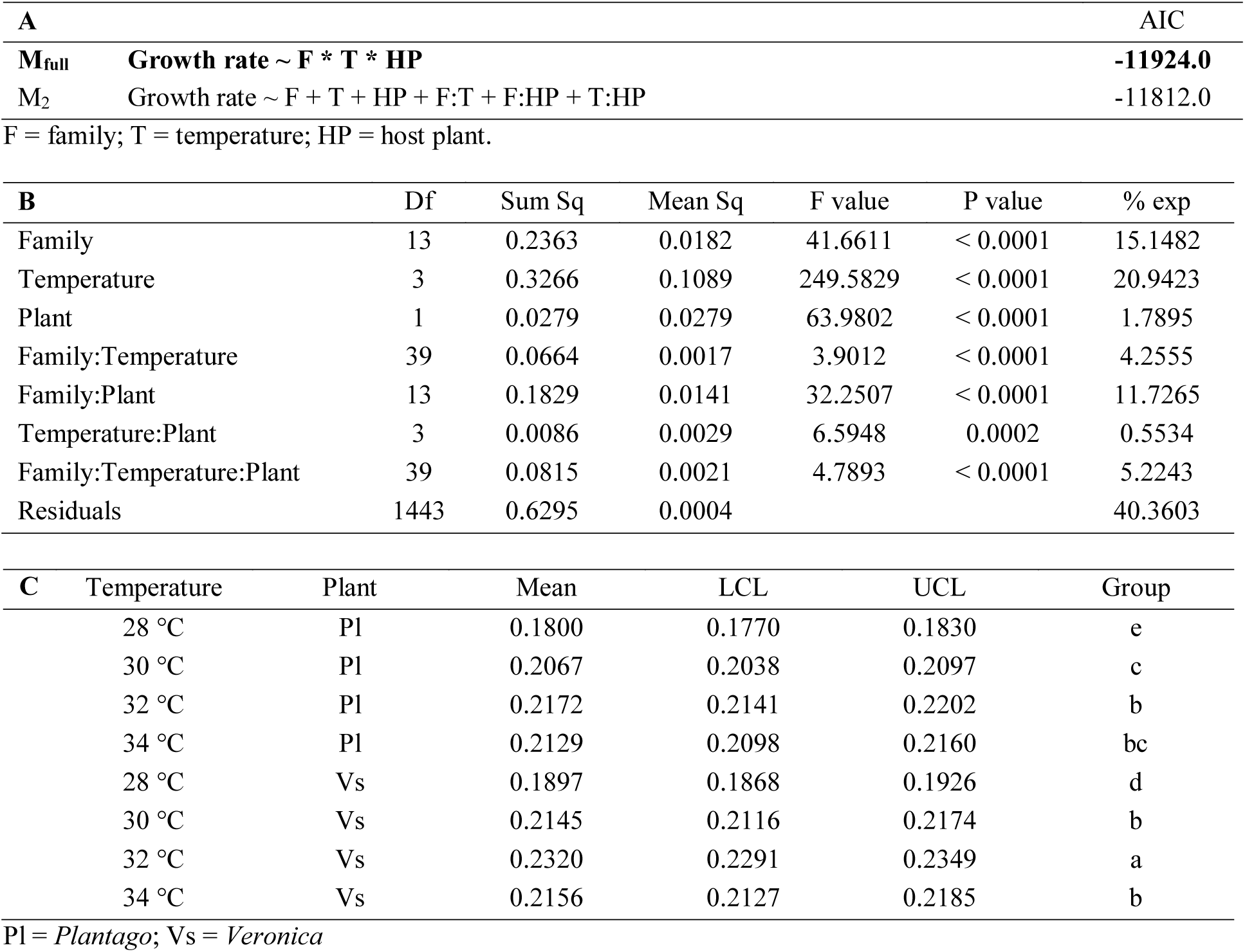
Linear model for individual growth rates (related to Figure 2B in the main text). **A**) Minimum adequate model (in bold) was obtained using the *step()* function, starting from the full model. **B**) Anova table for the minimum adequate model. **C**) Model-estimated marginal means, as well as upper and lower confidence limits, for all experimental treatments. Significant differences experimental groups (Tukey’s HSD, α = 0.05), averaged over the families, were obtained using the package *emmeans* and are indicated by different letters.

**Table S4:** Linear model for individual fat content (related to Figure 2B in the main text). **A**) Minimum adequate model (in bold) was obtained using the *step()* function, starting from the full model. **B**) Anova table for the minimum adequate model. **C**) Model-estimated marginal means, as well as upper and lower confidence limits, for all experimental treatments. Significant differences experimental groups (Tukey’s HSD, α = 0.05), averaged over the families, were obtained using the package *emmeans* and are indicated by different letters.

| **A** |  | AIC |
| --- | --- | --- |
| **M<sub>full</sub>** | **Fat content ~ F \* T \* HP** | **184.1** |
| M<sub>2</sub> | Fat content ~ F + T + HP + F:T + F:HP + T:HP | 236.5 |

F = family; T = temperature; HP = host plant.
| **B** | Df | Sum Sq | Mean Sq | F value | P value | % exp |
| --- | --- | --- | --- | --- | --- | --- |
| Family | 13 | 175.3395 | 13.4877 | 12.1996 | < 0.0001 | 11.4766 |
| Temperature | 3 | 95.9078 | 31.9693 | 28.9162 | < 0.0001 | 6.2775 |
| Plant | 1 | 33.0864 | 33.0864 | 29.9267 | < 0.0001 | 2.1656 |
| Family:Temperature | 39 | 204.0132 | 5.2311 | 4.7315 | < 0.0001 | 13.3534 |
| Family:Plant | 13 | 102.8951 | 7.9150 | 7.1591 | < 0.0001 | 6.7349 |
| Temperature:Plant | 3 | 16.7780 | 5.5927 | 5.0586 | 0.0018 | 1.0982 |
| Family:Temperature:Plant | 39 | 134.7115 | 3.4541 | 3.1243 | < 0.0001 | 8.8174 |
| Residuals | 692 | 765.0626 | 1.1056 |  |  | 50.0763 |

| **C** | Temperature | Plant | Mean | LCL | UCL | Group |
| --- | --- | --- | --- | --- | --- | --- |
|  | 28 °C | Pl | 7.0044 | 6.8002 | 7.2086 | a |
|  | 30 °C | Pl | 7.1823 | 6.9734 | 7.3912 | a |
|  | 32 °C | Pl | 7.6754 | 7.4670 | 7.8838 | b |
|  | 34 °C | Pl | 7.8885 | 7.6758 | 8.1012 | b |
|  | 28 °C | Vs | 6.7918 | 6.5859 | 6.9977 | a |
|  | 30 °C | Vs | 6.7932 | 6.5857 | 7.0007 | a |
|  | 32 °C | Vs | 6.7812 | 6.5715 | 6.9909 | a |
|  | 34 °C | Vs | 7.7098 | 7.5061 | 7.9136 | b |

Pl = *Plantago*; Vs = *Veronica*

**Table S5:**
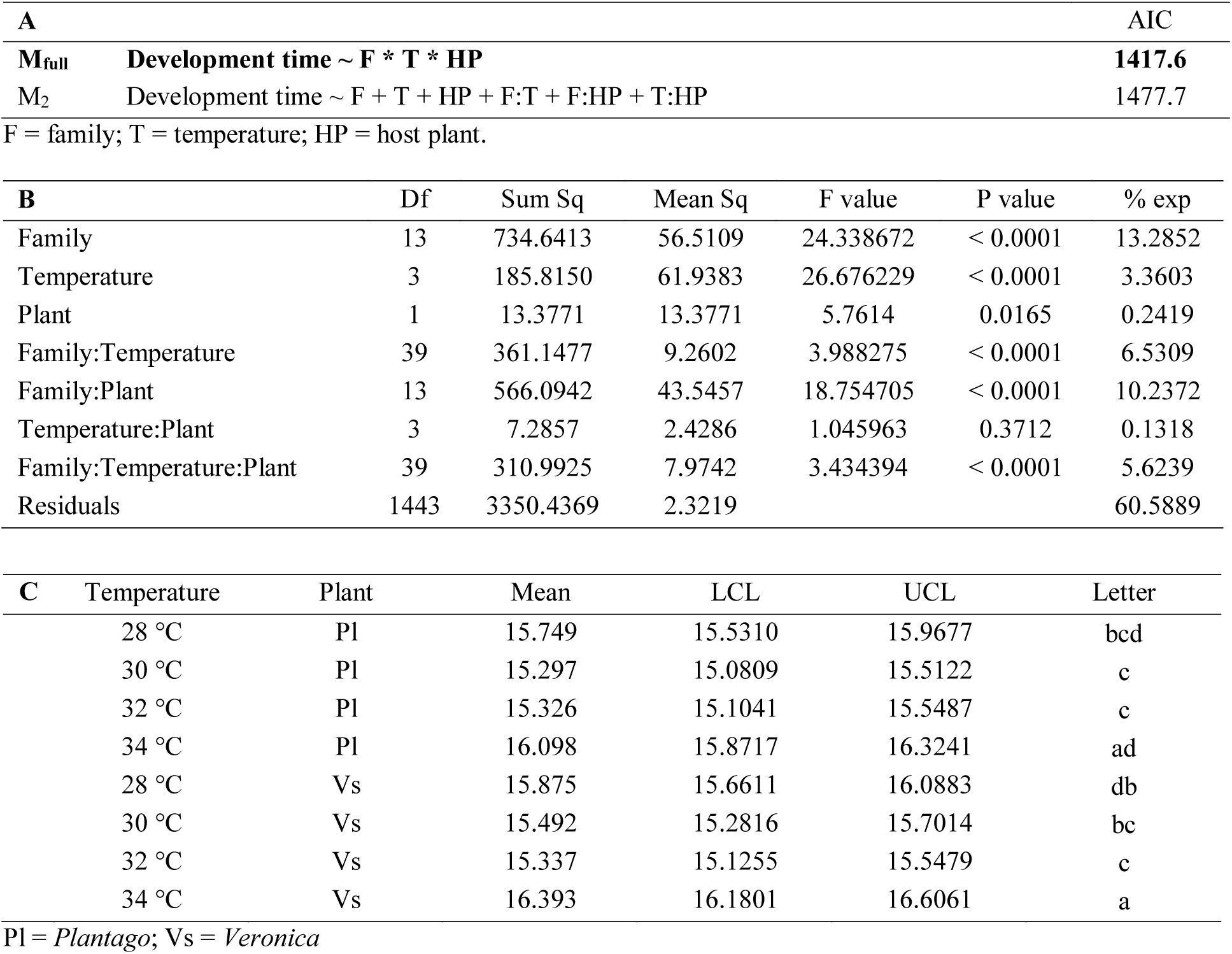
Linear model for individual development time (related to Figure S1A in the supplementary materials). **A**) Minimum adequate model (in bold) was obtained using the step() function, starting from the full model. **B**) Anova table for the minimum adequate model. **C**) Model-estimated marginal means, as well as upper and lower confidence limits, for all experimental treatments. Significant differences experimental groups (Tukey’s HSD, α = 0.05), averaged over the families, were obtained using the package emmeans and are indicated by different letters.

**Table S6:** Linear model for individual diapause mass (related to Figure S1B in the supplementary materials). **A**) Minimum adequate model (in bold) was obtained using the *step()* function, starting from the full model. **B**) Anova table for the minimum adequate model. **C**) Model-estimated marginal means, as well as upper and lower confidence limits, for all experimental treatments. Significant differences experimental groups (Tukey’s HSD, α = 0.05), averaged over the families, were obtained using the package *emmeans* and are indicated by different letters.

| A |  |  |  |  |  | AIC |
| --- | --- | --- | --- | --- | --- | --- |
| M<sub>full</sub> | Diapause mass ~ F * T * HP |  |  |  |  | 2891.9 |
| M<sub>2</sub> | Diapause mass ~ F + T + HP + F:T + F:HP + T:HP |  |  |  |  | 2938.8 |
| F = family; T = temperature; HP = host plant. |  |  |  |  |  |  |
| B | Df | Sum Sq | Mean Sq | F value | P value | % exp |
| Family | 13 | 3564.6910 | 274.2070 | 45.7602 | < 0.0001 | 19.4455 |
| Temperature | 3 | 3797.6067 | 1265.8689 | 211.2507 | < 0.0001 | 20.7160 |
| Plant | 1 | 169.6805 | 169.6805 | 28.3166 | < 0.0001 | 0.9256 |
| Family:Temperature | 39 | 758.5425 | 19.4498 | 3.2458 | < 0.0001 | 4.1379 |
| Family:Plant | 13 | 608.4506 | 46.8039 | 7.8107 | < 0.0001 | 3.3191 |
| Temperature:Plant | 3 | 63.2488 | 21.0829 | 3.5184 | 0.0146 | 0.3450 |
| Family:Temperature:Plant | 39 | 722.6794 | 18.5302 | 3.0924 | < 0.0001 | 3.9422 |
| Residuals | 1443 | 8646.8305 | 5.9923 |  |  | 47.1687 |
| C | Temperature | Plant | Mean | LCL | UCL | Letter |
|  | 28 °C | Pl | 6.8730 | 6.5222 | 7.2238 | e |
|  | 30 °C | Pl | 9.0712 | 8.7248 | 9.4176 | d |
|  | 32 °C | Pl | 10.0496 | 9.6925 | 10.4067 | bc |
|  | 34 °C | Pl | 10.6042 | 10.2408 | 10.9677 | bf |
|  | 28 °C | Vs | 7.1585 | 6.8154 | 7.5016 | e |
|  | 30 °C | Vs | 9.5695 | 9.2323 | 9.9067 | cd |
|  | 32 °C | Vs | 11.3837 | 11.0444 | 11.7230 | a |
|  | 34 °C | Vs | 11.2182 | 10.8760 | 11.5603 | af |

Pl = *Plantago*; Vs = *Veronica*

